# neoANT-HILL: an integrated tool for identification of potential neoantigens

**DOI:** 10.1101/603670

**Authors:** Ana Carolina M F Coelho, André L Fonseca, Danilo L Martins, Lucas M da Cunha, Paulo B R Lins, Sandro J de Souza

**Affiliations:** Bioinformatics Multidisciplinary Enviroment (BioME), Institute Metropolis Digital, Federal University of Rio Grande do Norte, UFRN, Brazil; PhD Program in Bioinformatics, UFRN, Natal, Brazil; Brain Institute, Federal University of Rio Grande do Norte, UFRN, Brazil

**Keywords:** neoantigens, cancer, immunogenomic analysis

## Abstract

Cancer neoantigens have attracted great interest in immunotherapy due to their ability to elicit antitumoral immune responses. These antigens are formed due to somatic mutations in the cancer genome that result in alterations of the original protein. Although current technological advances in neoantigen identification, it remains a challenging and a large number of false-positive continue to exist. In the current work, we present neoANT-HILL, an automatized user-friendly tool that integrates several immunogenomic analysis to improve neoantigens detection from NGS data. The program input can be a file with somatic mutations called and/or RNA-seq data. Our tool was applied on somatic mutations of melanoma dataset from TCGA and found that neoANT-HILL was able to predicted potential neoantigens. The software is available on github at https://github.com/neoanthill/neoANT-HILL.

## 1 Introduction

Recent studies have demonstrated that T cells can recognize tumor-specific antigens that bound to the human leukocyte antigens (HLA) molecules at the surface of tumor cells (Efremova *et al.*, 2017; Kato *et al*., 2018). During tumor progression, accumulating somatic mutations in the tumor genome can affect protein-coding genes and result in mutated peptides (Efremova *et al*., 2017). These mutated peptides, which are present in the malignant cells but not in the normal cells, may act as neoantigens and trigger T-cell responses due to the lack of thymic elimination of autoreactive T-cells (central tolerance) (Snyder *et al*., 2014; Bailey *et al*., 2016; Riaz *et al*., 2016). As result, these neoantigens appear to represent ideal targets attracting great interest for cancer immunotherapeutic strategies, including therapeutic vaccines and engineered T cells (Lu; Robbins, 2016; Efremova *et al*., 2017).

In the last years, advances in next-generation sequencing have provided an accessible way to generate patient-specific data, which allows the prediction of tumor neoantigens in a rapid and comprehensive manner (Liu; Mardis, 2017). Several approaches have been developed, such as pVAC-Seq (Hundal *et al.,* 2016), MuPeXI (Bjerregaard *et al.,* 2017), TIminer (Tappeiner *et al*., 2017) and TSNAD (Zhou *et al*., 2017), which performs prediction of potential neoantigens produced by non-synonymous mutations. However, none of these proposed tools considers tumor transcriptome sequencing data (RNA-seq) for identifying somatic mutations. Moreover, only one of these tools provides quantification of the fraction of tumor-infiltrating immune cells types.

Here we are presenting a versatile tool with a graphical user interface (GUI), called neoANT-HILL, designed to identify potential neoantigens arising from cancer somatic missense mutations, frameshift and small indels. neoANT-HILL integrates several complementary features to prioritizing mutant peptides based on predicted binding affinity and mRNA expression level levels. We used datasets from GEUVADIS RNA sequencing project (Lappalainen *et al*., 2013) to demonstrate that RNA-seq is also a potential source of mutation detection. Finally, we applied our pipeline on a large melanoma cohort from The Cancer Genome Atlas (Weinstein *et al*., 2013) to demonstrate its utility in predicting and suggesting potential neoantigens.

## 2 Material and Methods

### RNA-seq data processing

We obtained RNA-seq samples (*n=15*) from the GEUVADIS RNA sequencing project to identifying frameshift, indels and point mutations. Raw RNA-seq reads were mapped with STAR aligner (version 2.6.0) (Dobin, *et al*., 2013) in two-pass mapping protocol against the human reference genome version GRCh37. Mapped reads were processed according to GATK best practices (DePristo *et al*., 2011; Van der Auwera *et al*., 2013). Mutect2 (Cibulskis *et al*., 2013) was used to identifies frameshift, missense mutations and small indels. We limited our analysis to variants that presented a read depth (DP) >= 10 and were supported by at least five reads. Our results were validated by comparing the corresponding genotype provided by 1000 Genomes Project Consortium (1KG) (1000 Genomes Project Consortium *et al*., 2015). Bcftools *isec* was used to determine the intersection between the variant sets.

### Melanoma dataset

We applied neoANT-HILL on a large melanoma cohort (SKCM, *n* = 466) obtained from The Genome Cancer Atlas (TCGA) to identifying potential neoantigens. Expression levels (FPKM) from corresponding samples were also obtained. All missense mutations, frameshift and small indels were extracted. The mutant sequences were inferred and reported with its corresponding wild-type sequence. Our analysis was limited to HLA class I molecules. We used a set of HLA molecules that were within the most frequent alleles collected in the 1KG Project, including HLA-A*02:01, HLA-A*11:01, HLA-A*24:02, HLA-B*07:02, HLA-B*15:01, HLA-C*06:02 and HLA-C*07:02. The binding affinity prediction was run using ANN algorithm (v. 4.0 *aka* NetMHC) provided by IEDB for lengths 8-, 9-, 10- and 11-mer. We selected mutant peptides which its matched normal peptide showed a predicted binding affinity >= 500 nM. We also consider only the mutant peptides with the lowest predicted IC50 per HLA allele to avoid overlapping candidates differing by the length.

## 3 Results

### neoANT-HILL overview and availability

neoANT-HILL is a user-friendly integrated tool for the identification of potential neoantigens that could be used in personalized immunotherapy (Figure 1). Our pipeline relies on VCF file (single-or multisample) or tumor transcriptome sequence data (RNA-seq) in which somatic mutation calling will be performed following GATK best practices with Mutect2. In the current implementation, neoANT-HILL supports VCF files generated using the human genome version GRCh37. Other human genome version must be converted to version GRCh37. A list of HLA alleles should also be provided. At first, the variants are properly annotated by snpEff (Cingolani *et al.*, 2012). The next step is identifying non-synonymous mutations (missense mutations, frameshift and indels) and infers the resulting mutant sequence. The protein sequence changes are inferred from the NCBI Reference Sequence database (RefSeq) (O’Leary *et al*., 2016). For frameshift mutations, the mutant sequence is inferred by translating the cDNA sequence. Each alteration is translated into a 21-mer sequence where the altered point is at the center. If the mutation is at the beginning or the end of the transcript, the mutant sequence is built by taking the 20 succeeding or preceding amino acids, respectively. The translated mutant sequences and its wild-type corresponding sequence are stored in a FASTA file.

**Figure 1.**
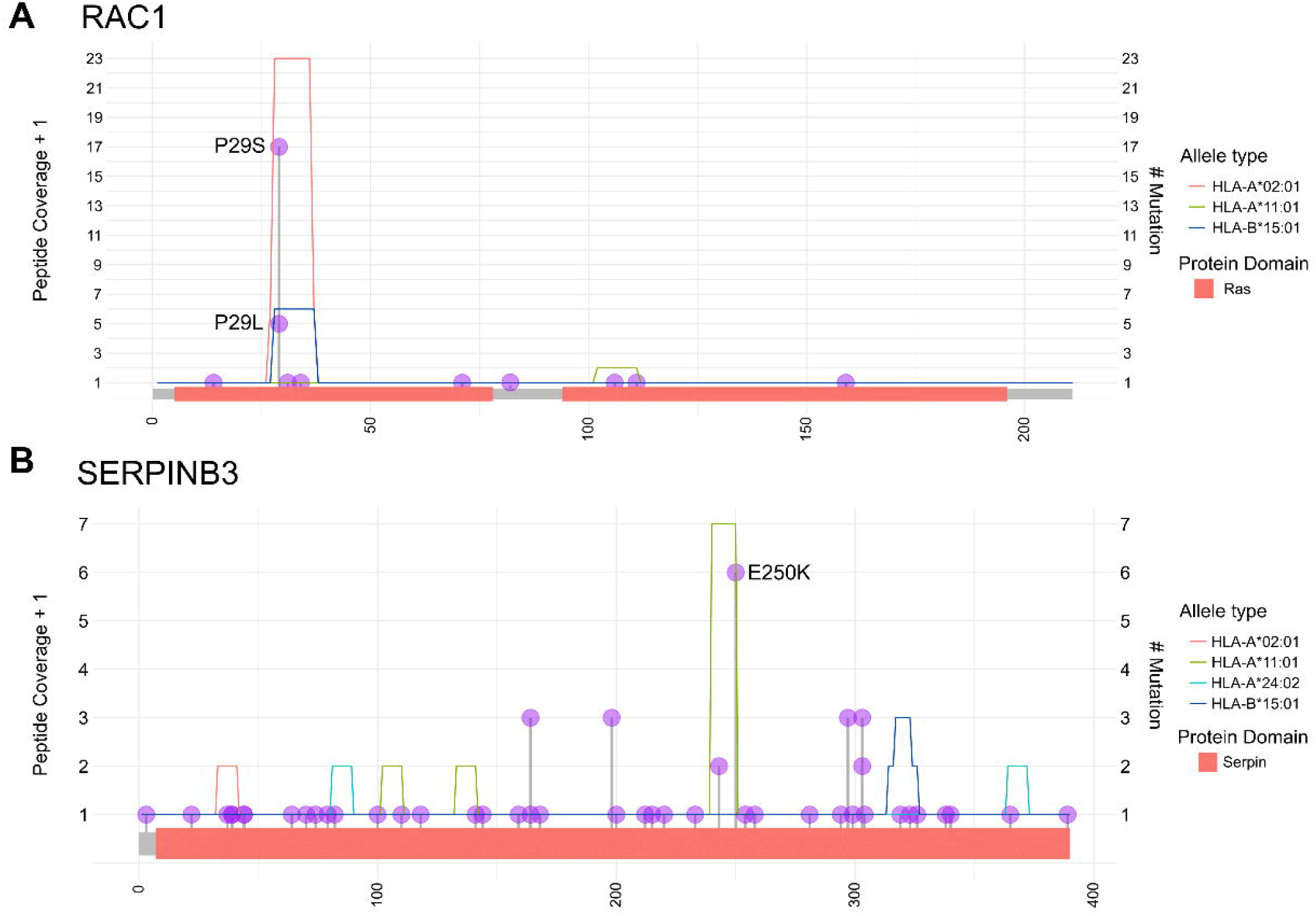
Overall workflow of neoANT-HILL.

The subsequent step is binding affinity prediction between the peptides and HLA alleles. neoANT-HILL supports seven HLA class I molecules algorithms provided by Immune Epitope Database (IEDB) (VITA *et al*., 2015), including NetMHC (v. 4.0) (Andreatta; Nielsen, 2016; Nielsen *et al*., 2003), NetMHCpan (v. 4.0) (Jurtz *et al.,* 2017), NetMHCcons (Karosiene *et al.,* 2012), NetMHCstabpan (Rasmussen *et al.,* 2016), PickPocket (Zhang; Lund; Nielsen, 2009), SMM (Peters; Sette, 2005) and SMMPMBEC (Kim *et al*., 2009) and MHCflurry (O’Donnell *et al*., 2018). Each peptide sequence is parsed with a sliding window metric. The algorithm also allows the prediction of binding affinity for HLA class II through four IEDB-algorithms NetMHCIIpan (v. 3.1) (Karosiene *et. al*, 2013), NN-align (Nielsen; Lund, 2009), SMM-align (Nielsen; Lundgaard; Lund, 2007) and Sturniolo (Sturniolo *et al*., 1999). It can be executed on parallel single or multi-sample using parallelization with the custom configured parameters. The binding affinities are predicted to both mutated and normal peptides. The differential agretopicity index (DAI) (Ghorani *et al*., 2018) is also reported, which represent the fold change between normal and mutated peptides binding affinities.

Moreover, if raw RNA-seq data is available (in FASTQ format), neoANT-HILL pipeline can perform complementary analyses. Our algorithm uses Optitype (Szolek *et al*., 2014) to infers class-I HLA molecules. The data can also be used to estimate gene and transcript level expression, which is reported in transcripts per million (TPM), using Kallisto (Bray *et al.*, 2016). Genes are considered to be expressed if they show an abundance level of at least 1 TPM. In addition, neoANT-HILL also offers the possibility of estimating quantitatively, via deconvolution, the relative fractions of tumor-infiltrating immune cell types through the use of quanTIseq (Finotello *et al*., 2017).

Our software was developed under a pre-built Docker image. The required dependencies are packaged up which simplifying the installation process and avoid possible incompatibilities between versions. It can be installed on Unix/Linux, MacOS, and Windows operating systems. neoANT-HILL was designed through a user graphical interface (Figure 2) implemented on Flask framework. As previously described, several analyses are supported and each one relies on different tools. Several scripts were implemented on Python (v. 2.7) to complete automating the execution of these single tools and data integration. The results of each analysis are stored in separate tabs on the sample-specific folder. They are shown in tabular or graphical forms that let the user manage the data based on their own selection criteria.

**Figure 2.**
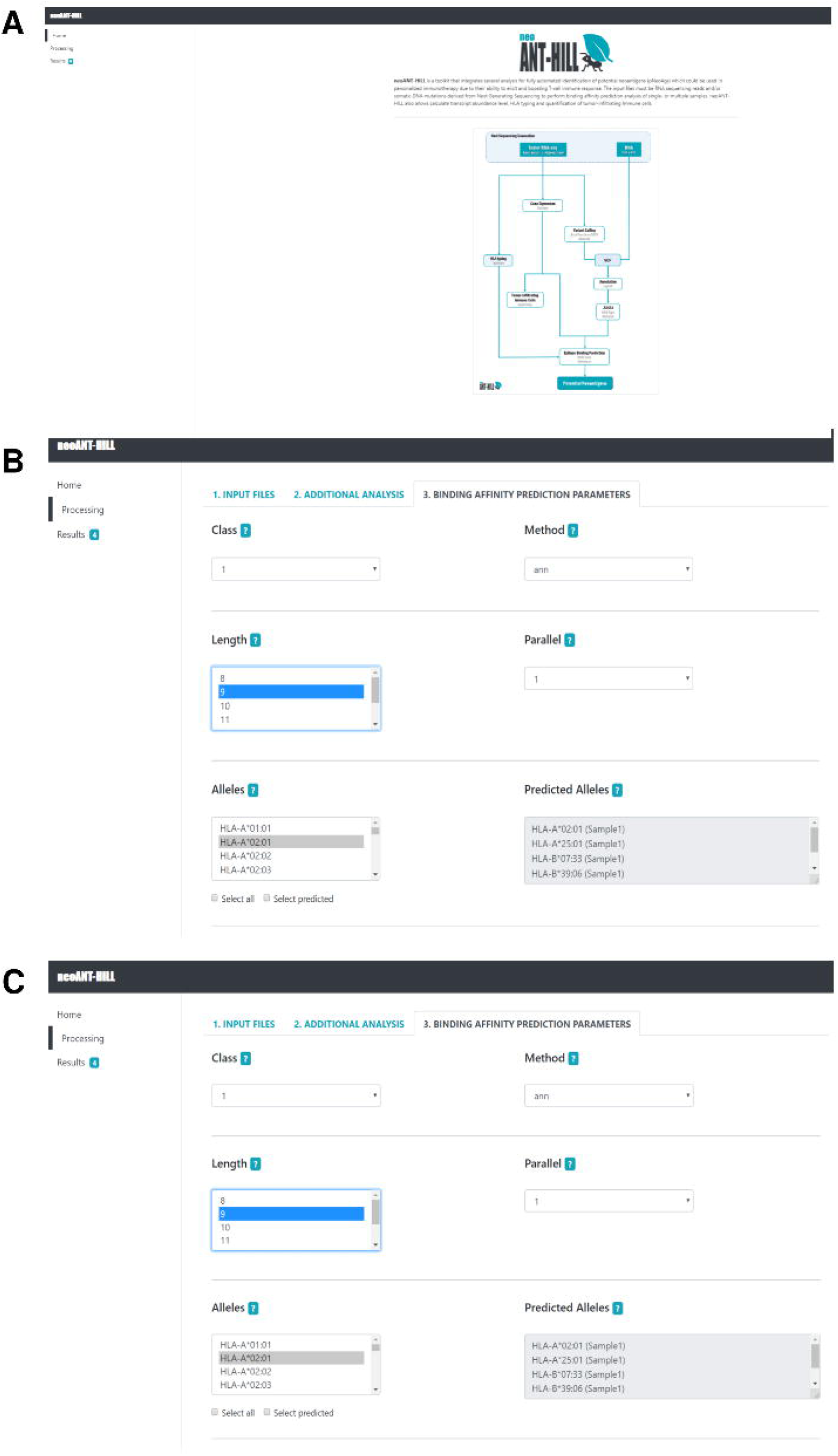
Screenshots of neoANT-HILL interface. **(A)** Main page of neoANT-HILL. **(B)** Processing tab for parameters selection to run binding prediction affinity. **(C)** Binding prediction results tab.

### Non-synonymous mutations identification on RNA-seq

We evaluate the utility of RNA-seq for identifying frameshift, indels and point mutations. We used samples from the GEUVADIS project since for those samples a reference genome was available through the 1K Genome project. Although these samples are not derived from tumor cells, the goal of these analysis was to benchmark the efficiency of our pipeline to detect somatic mutations from RNA-Seq data. Mutect2 was performed on tumor-only mode without distinction between somatic and germline variants. The overall called variants were then compared to the corresponding genotypes. We found that on average 72% of variants in coding regions detected by RNA-seq were confirmed by the genome sequencing (concordant calls) (Supplementary Table1). Variants in genes that are not expressed cannot be detected by RNA-seq. Mapping mismatches and RNA editing sites could partially explain discordant calls.

### Predicted potential neoantigens on melanoma

We found approximately 198,000 records of predicted mutant peptides in the SKCM dataset from the TCGA project. It is important to note that the large amount of mutant peptides is due to the high mutational burden of melanoma and the set of HLA alleles that was used to run the binding prediction. Moreover, these mutant peptides were classified as strong (IC50 < 50 nM), intermediate (IC50 >= 50 nM and < 250 nM) and weak binder (IC50 >= 250 nM and < 500 nM) (Supplementary Table 2). We decided to focus on expressed mutant peptides classified as strong binders to further evaluation as potential neoantigens.

We observed that the distribution of the majority of strong binders mutant peptides are private and unique, which demonstrates the intratumor heterogeneity. However, we observed that frequent mutations may be likely to generate recurrent mutant peptides (Table 1). For instance, a potential neoantigen (F**<u>S</u>**GEYIPTV), which was predicted to form a complex with HLA-A*02:01 allele, was found to be shared among 17 samples (3.65%). It was generated from the P29S mutation in gene RAC1. Another mutation (P29L) in the same gene was also related to form a recurrent potential neoantigen (F**<u>L</u>**GEYIPTV) that was found in 5 samples (1.07%). As another example, we can also highlight another potential shared neoantigen (LSMIVLLPN**<u>K</u>**) related to mutation E250K in the SERPINB3 gene (Figure 3B). It was found in 6 samples (1.29%) and it was likely to form a complex with the HLA-A*11:01 allele.

**Table 1.**
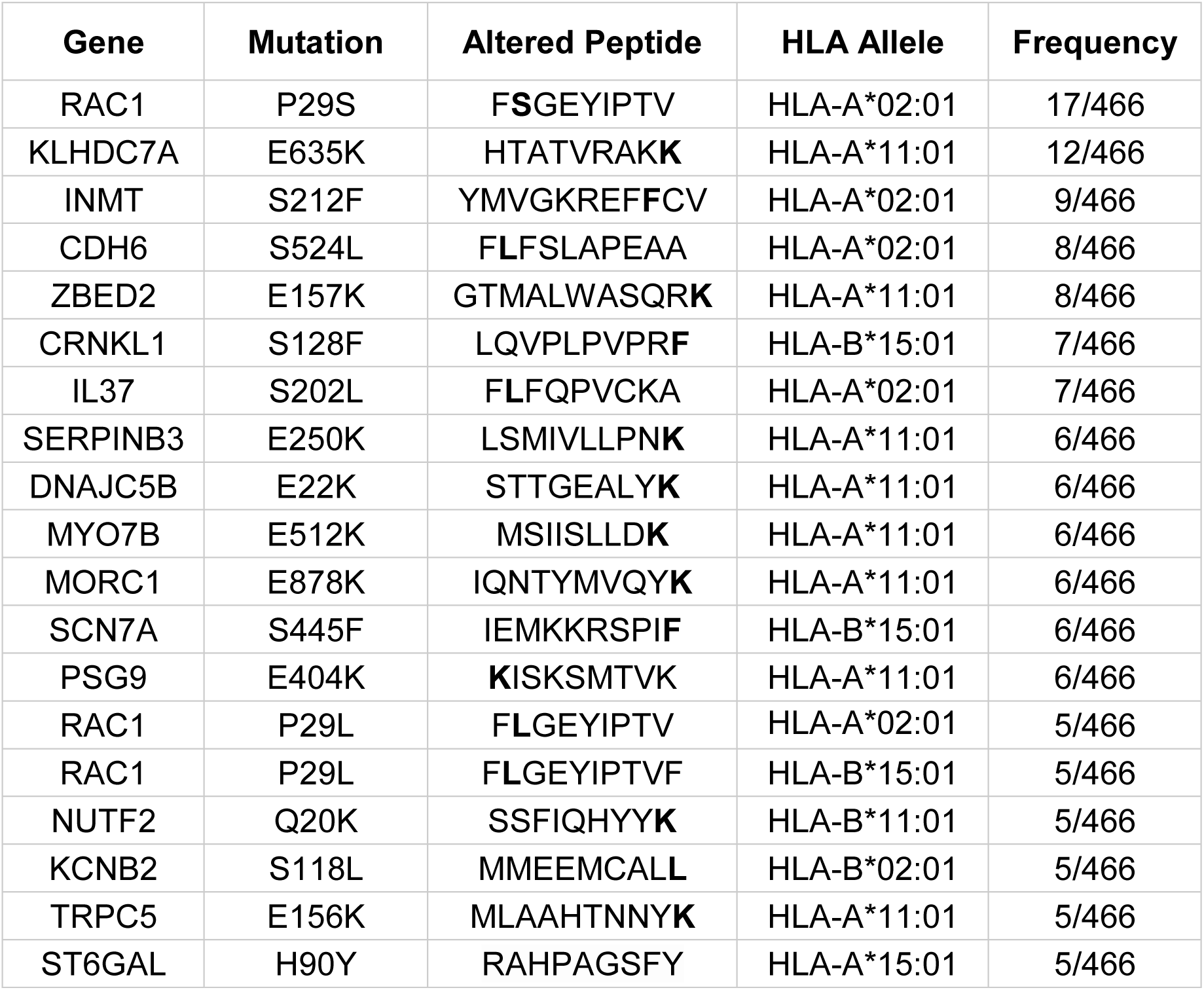
Top 20 potential shared neoantigens.

**Figure 3.**
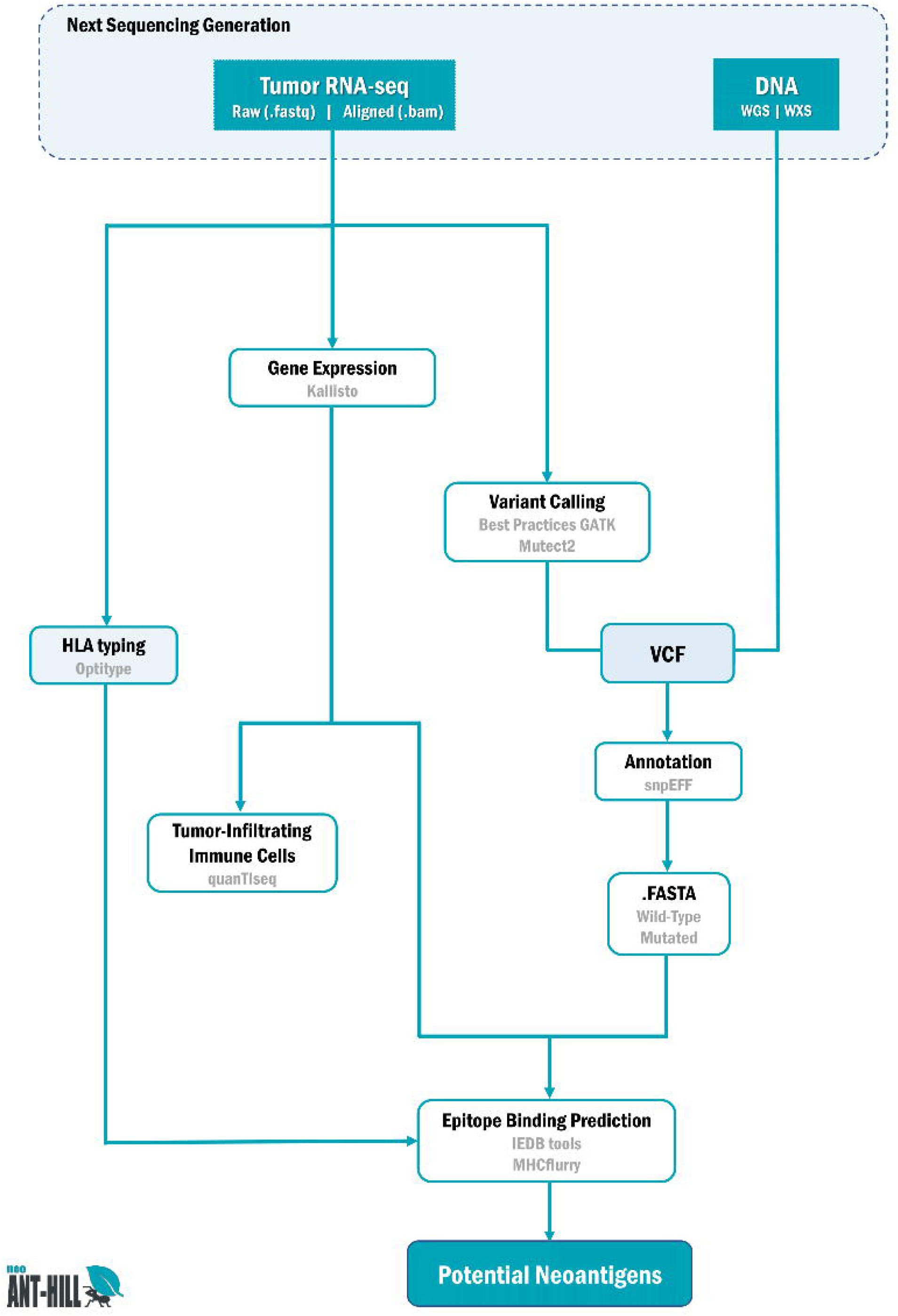
Distribution of recurrent missense mutations that generated potential shared neoantigens. (**A)** P29S and RAC1 gene generated recurrent strong binders mutant peptides to HLA-A*02:01 and P29L generated strong binders that could respond to HLA-A*02:01 or HLA-A*11:01, depending on peptide length **(B)** E250K in SERPINB3 gene generate a recurrent potential neoantigen that binds to HLA-A*11:01.

We also observed that overlapping sequences of different lengths have shown more stable binding to different alleles (Supplementary Table 3). For instance, the previous mentioned peptide FLGEYIPTV, related to P29L RAC1, was likely to strongly bind to HLA-A*02:01. While the decameter peptide <u>FLGEYIPTV</u>F, which is also related to the same mutation, have been show to respond to HLA-A*11:01 (Figure 3A). Similarly, the peptide AQIEASLSV, from R414Q HHATL, have been shown to strongly respond to HLA-A*02:01, while L<u>AQIEASLSV</u> bound to HLA-A*15:01.

## 4 Discussion

Cancer immunotherapy is rapidly advancing due to the progress in the understanding of the interaction between cancer and immune cells. Neoantigens are attractive candidates because these peptides can be used to design a personalized, efficient and safer cancer immunotherapy option (Guo *et al.*, 2018). However, accurate prediction of neoantigens remains a challenge due to multiple factors such as antigen processing, HLA binding affinity, amino acid composition and expression level of the mutant peptide that must be considered (Ghorani *et al.* 2017). Here we presented neoANT-HILL which covers and integrates many of these specific sub-tasks. Our tool also has the ability to explore the versatility of RNA sequencing including variant calling, in abscence of DNA sequencing data, gene expression level, inferencing of HLA type and profiling tumor-infiltrating immune cells.

Although calling variants from RNA-Seq data has been shown to be more challenging, it is a interesting alternative for genome sequencing and a large amount of tumor RNA-seq samples do not have normal matched data (Piskol; Ramaswami; Li, 2013; Coudray *et al.,* 2018). We applied the variant calling pipeline on RNA-seq data from GEUVADIS and we demonstrated the feasibility of variants detection with remarkable precision. In addition, another complementary step, which is explored by neoANT-HILL, is quantifying tumor-infiltrating immune cells from RNA-seq data. It has been demonstrated that the evaluation of tumor-infiltrating lymphocytes (TILs) provides prognostic value and potential predictive information of response to immunotherapy (Gooden *et al*., 2011; Althobiti *et al*., 2018). In comparison to the previously proposed tools, they usually considers RNA-seq data to estimate gene expression level or HLA typing. Only TIminer provides the option of quantifying the tumor-infiltrating immune cells through gene set enrichment analysis (GSEA).

We have also used melanoma dataset from TCGA to demonstrate the utility of neoANT-HILL in identifying potential neoantigens. We found several predicted patient specific and shared neoantigen candidates. The use of non-patient specific HLA alleles in this analysis may have generated false positive potential neoantigens. We observed that recurrent mutations in RAC1 and SERPINB3 genes are likely to form potential neoantigens. RAC1 P29S have been described as a candidate biomarker for treatment with anti-PD1 or anti-PD-L1 antibodies (Vu *et al*., 2015). Mutations in SERPINB3 have also been related to response to immunotherapy (Riaz *et al*. 2016). Therefore, our results suggests that screening these neoantigens can be used as predictive biomarkers for immune responses and potential targets for immunotherapies.

Our tool provides completely integrated analyses to predicting potential neoantigens candidates. neoANT-HILL is available through a user-friendly graphical interface which enables its usage by users without an advanced programming background. However, neoANT-HILL still lacks some features that must be taken into consideration in future updates such as detection of mutations that arise from gene fusion, inference of HLA-class II and evaluation of similarity to known epitopes.

### Software availability

neoANT-HILL is hosted publicly on GitHub at https://github.com/neoanthill/neoANT-HILL and the user documentation is also available on this page.

## Data availability

The RNA-Seq dataset from Geuvadis RNA sequencing project were downloaded from the ArrayExpress database (http://www.ebi.ac.uk/arrayexpress/) and this data can be accessed under the accession number <u>E-GEUV-1</u>. We used the individuals named NA12812, NA12749, NA20510, NA19119, NA19204, NA18498, NA12489, NA20752, NA18517, NA11992, NA19144, NA20759, NA19137, NA19257 and NA12006. The corresponding genotyping data (Phase I) were downloaded from the data portal of the 1KG Project (http://www.internationalgenome.org/). The melanoma TCGA mutation and expression data were obtained from cBIO portal by using the CGDS-R package.

## Author Contributions

ACMFC, DLM and PRBL designed and carried out the implementation of the computational pipeline. LMC contributed to design the computational pipeline. ACMFC and ALF analyzed the data. ACMFC wrote the manuscript in consultation with SJS. SJS supervised the project.

## REFERENCES

1. Efremova, M. et al. Neoantigens generated by individual mutations and their role in cancer immunity and immunotherapy. Frontiers in immunology, v. 8, p. 1679, 2017. doi:10.3389/fimmu.2017.01679

2. Kato, T. et al. Effective screening of T cells recognizing neoantigens and construction of T-cell receptor-engineered T cells. Oncotarget, v. 9, n. 13, p. 11009, 2018

3. Snyder, A. et al. Genetic basis for clinical response to CTLA-4 blockade in melanoma. New England Journal of Medicine, v. 371, n. 23, p. 2189–2199, 2014.

4. Bailey, P. et al. Exploiting the neoantigen landscape for immunotherapy of pancreatic ductal adenocarcinoma. Scientific reports, v. 6, p. 35848, 2016.

5. Riaz, N. et al. The role of neoantigens in response to immune checkpoint blockade. International immunology, v. 28, n. 8, p. 411–419, 2016.

6. Lu, Y.-C.; Robbins, P. F. Cancer immunotherapy targeting neoantigens. In: Seminars in immunology. Academic Press, 2016. p. 22–27.

7. Liu, X. S.; Mardis, E. R. Applications of immunogenomics to cancer. Cell, v. 168, n. 4, p. 600–612, 2017.

8. Hundal, J. et al. pVAC-Seq: A genome-guided in silico approach to identifying tumor neoantigens. Genome medicine, v. 8, n. 1, p. 11, 2016.

9. Bjerregaard, A.-M. et al. MuPeXI: prediction of neo-epitopes from tumor sequencing data. Cancer Immunology, Immunotherapy, v. 66, n. 9, p. 1123–1130, 2017.

10. Tappeiner, E. et al. TIminer: NGS data mining pipeline for cancer immunology and immunotherapy. Bioinformatics, v. 33, n. 19, p. 3140–3141, 2017.

11. Zhou, Z. et al. TSNAD: an integrated software for cancer somatic mutation and tumour-specific neoantigen detection. Royal Society open science, v. 4, n. 4, p. 170050, 2017.

12. Lappalainen, T. et al. Transcriptome and genome sequencing uncovers functional variation in humans. Nature, v. 501, n. 7468, p. 506, 2013.

13. Weinstein, J. N. et al. The cancer genome atlas pan-cancer analysis project. Nature genetics, v. 45, n. 10, p. 1113, 2013.

14. Dobin, A. et al. STAR: ultrafast universal RNA-seq aligner. Bioinformatics, v. 29, n. 1, p. 15–21, 2013.

15. Depristo, M. A. et al. A framework for variation discovery and genotyping using next-generation DNA sequencing data. Nature genetics, v. 43, n. 5, p. 491, 2011.

16. Van Der Auwera, G. A. et al. From FastQ data to high-confidence variant calls: the genome analysis toolkit best practices pipeline. Current protocols in bioinformatics, v. 43, n. 1, p. 11.10.1-11.10. 33, 2013.

17. Cibulskis, K. et al. Sensitive detection of somatic point mutations in impure and heterogeneous cancer samples. Nature biotechnology, v. 31, n. 3, p. 213, 2013.

18. 1000 GENOMES PROJECT CONSORTIUM et al. A global reference for human genetic variation. Nature, v. 526, n. 7571, p. 68, 2015.

19. Cingolani, P. et al. A program for annotating and predicting the effects of single nucleotide polymorphisms, SnpEff: SNPs in the genome of Drosophila melanogaster strain w1118; iso-2; iso-3. Fly, v. 6, n. 2, p. 80–92, 2012.

20. O’Leary, N. A. et al. Reference sequence (RefSeq) database at NCBI: current status, taxonomic expansion, and functional annotation. Nucleic acids research, v. 44, n. D1, p. D733–D745, 2015.

21. VITA Randi et al. The immune epitope database (IEDB) 3.0. Nucleic acids research, v. 43, n. D1, p. D405-D412, 2014.

22. Andreatta, M.; Nielsen, M. Gapped sequence alignment using artificial neural networks: application to the MHC class I system. Bioinformatics, v. 32, n. 4, p. 511–517, 2015.

23. Nielsen, M. et al. Reliable prediction of T-cell epitopes using neural networks with novel sequence representations. Protein Science, v. 12, n. 5, p. 1007–1017, 2003.

24. Jurtz, v. et al. NetMHCpan-4.0: Improved peptide–MHC class I interaction predictions integrating eluted ligand and peptide binding affinity data. The Journal of Immunology, v. 199, n. 9, p. 3360–3368, 2017.

25. Karosiene, E. et al. NetMHCcons: a consensus method for the major histocompatibility complex class I predictions. Immunogenetics, v. 64, n. 3, p. 177–186, 2012.

26. Rasmussen, M. et al. Pan-Specific Prediction of Peptide–MHC Class I Complex Stability, a Correlate of T Cell Immunogenicity. The Journal of Immunology, v. 197, n. 4, p. 1517–1524, 2016.

27. Zhang, H., Lund, O., Nielsen, M. The PickPocket method for predicting binding specificities for receptors based on receptor pocket similarities: application to MHC-peptide binding. Bioinformatics, v. 25, n. 10, p. 1293–1299, 2009.

28. Peters, B., Sette, A. Generating quantitative models describing the sequence specificity of biological processes with the stabilized matrix method. BMC bioinformatics, v. 6, n. 1, p. 132, 2005.

29. Kim, Y. et al. Derivation of an amino acid similarity matrix for peptide: MHC binding and its application as a Bayesian prior. BMC bioinformatics, v. 10, n. 1, p. 394, 2009.

30. O’Donnell, T. J. et al. MHCflurry: open-source class I MHC binding affinity prediction. Cell systems, v. 7, n. 1, p. 129-132. e4, 2018.

31. Karosiene, E. et al. NetMHCIIpan-3. 0, a common pan-specific MHC class II prediction method including all three human MHC class II isotypes, HLA-DR, HLA-DP and HLA-DQ. Immunogenetics, v. 65, n. 10, p. 711–724, 2013.

32. Nielsen, M., Lund, O. NN-align. An artificial neural network-based alignment algorithm for MHC class II peptide binding prediction. BMC bioinformatics, v. 10, n. 1, p. 296, 2009.

33. Nielsen, M., Lundegaard, C., Lund, O. Prediction of MHC class II binding affinity using SMM-align, a novel stabilization matrix alignment method. BMC bioinformatics, v. 8, n. 1, p. 238, 2007.

34. Sturniolo, T. et al. Generation of tissue-specific and promiscuous HLA ligand databases using DNA microarrays and virtual HLA class II matrices. Nature biotechnology, v. 17, n. 6, p. 555, 1999.

35. Ghorani, E. et al. Differential binding affinity of mutated peptides for MHC class I is a predictor of survival in advanced lung cancer and melanoma. Annals of Oncology, v. 29, n. 1, p. 271–279, 2017.

36. Szolek, A. et al. OptiType: precision HLA typing from next-generation sequencing data. Bioinformatics, v. 30, n. 23, p. 3310–3316, 2014.

37. Bray, N. L. et al. Near-optimal probabilistic RNA-seq quantification. Nature biotechnology, v. 34, n. 5, p. 525, 2016.

38. Finotello, F. et al. Molecular and pharmacological modulators of the tumor immune contexture revealed by deconvolution of RNA-seq data. bioRxiv, p. 223180, 2018.

39. Guo, Y., Lei, K., Tang, L. Neoantigen vaccine delivery for personalized anticancer immunotherapy. Frontiers in immunology, v. 9, p. 1499, 2018.

40. Piskol, R., Ramaswami, G., Li, J. B. Reliable identification of genomic variants from RNA-seq data. The American Journal of Human Genetics, v. 93, n. 4, p. 641–651, 2013.

41. Coudray, A. et al. Detection and benchmarking of somatic mutations in cancer genomes using RNA-seq data. PeerJ, v. 6, p. e5362, 2018.

42. Coudray, A. et al. Detection and benchmarking of somatic mutations in cancer genomes using RNA-seq data. PeerJ, v. 6, p. e5362, 2018.

43. Althobiti, M. et al. Heterogeneity of tumour-infiltrating lymphocytes in breast cancer and its prognostic significance. Histopathology, v. 73, n. 6, p. 887–896, 2018.

44. Vu, H. L. et al. RAC 1 P29S regulates PD-L1 expression in melanoma. Pigment cell & melanoma research, v. 28, n. 5, p. 590–598, 2015.

45. Riaz, N. et al. Recurrent SERPINB3 and SERPINB4 mutations in patients who respond to anti-CTLA4 immunotherapy. Nature genetics, v. 48, n. 11, p. 1327, 2016.

